# Toxicokinetic Characterization of the Inter-Species Differences in 6PPD-Quinone Toxicity Across Seven Fish Species: Metabolite Identification and Semi-Quantification

**DOI:** 10.1101/2023.08.18.553920

**Authors:** David Montgomery, Xiaowen Ji, Jenna Cantin, Danielle Philibert, Garrett Foster, Summer Selinger, Niteesh Jain, Justin Miller, Jenifer McIntyre, Benjamin de Jourdan, Steve Wiseman, Markus Hecker, Markus Brinkmann

## Abstract

N-(1,3-Dimethylbutyl)-N’-Phenyl-P-Phenylenediamine-Quinone (6PPD-Q) is a recently identified contaminant that originates from the oxidation of the tire anti-degradant 6PPD. 6PPD-Q is acutely toxic to select salmonids at environmentally relevant concentrations, while other fish species display tolerance to concentrations surpassing those measured in the environment. The reasons for these marked differences in sensitivity are presently unknown. The objective of this research was to explore potential toxicokinetic drivers of species sensitivity by characterizing biliary metabolites of 6PPD-Q in sensitive and tolerant fishes. For the first time, we identified an *O*-glucuronide metabolite of 6PPD-Q using high-resolution mass spectrometry. The semi-quantified levels of this metabolite in tolerant species or life stages, including white sturgeon (*Acipenser transmontanus*), chinook salmon (*Oncorhynchus tshawytscha*), westslope cutthroat trout (*Oncorhynchus clarkia lewisi*) and non-fry life stages of Atlantic salmon (*Salmo salar*), were greater than those in sensitive species, including coho salmon (*Oncorhynchus kisutch*), brook trout (*Salvelinus fontinalis*), and rainbow trout (*Oncorhynchus mykiss*), suggesting that tolerant species might more effectively detoxify 6PPD-Q. Thus, we hypothesize that differences in species sensitivity are a result of differences in basal expression of biotransformation enzyme across various fish species. Moreover, the semi-quantification of 6PPD-Q metabolites in bile extracted from wild-caught fish might be a useful biomarker of exposure to 6PPD-Q, thereby being invaluable to environmental monitoring and risk assessment.

## INTRODUCTION

N-(1,3-Dimethylbutyl)-N’-Phenyl-P-Phenylenediamine-Quinone (6PPD-Q), an oxidation product of the tire anti-degradant 6PPD, is a novel contaminant of great global concern due to its high ecotoxicity.^1-12^ 6PPD belongs to the class of *p*-phenylenediamine compounds, which are aromatic amine antioxidants that function to increase the lifespan of pneumatic rubber tires and were invented in 1845.^2,3^ In the environment, 6PPD rapidly oxidizes into its di-ketone (or quinone) product, 6PPD-Q.^1^ This oxidation product has recently been shown to be highly toxic to a variety of salmonid species, thus constituting a great global concern due to its ubiquitous nature.^1, 4-12^ 6PPD-Q is transported into nearby waterways with roadway runoff during rainfall events, and maximum surface water concentrations of 0.6 to 2.3 μg/L have been reported across North America (e.g., Seattle, WA, USA; Saskatoon, SK, Canada; and Toronto, ON, Canada).^4-6^The toxicity of 6PPD-Q was first discovered by Tian et al. ^5,6^ in coho salmon (*Oncorhynchus kisutch*; abbreviated: CS). This species showed mass mortalities after acute exposure to environmentally relevant concentrations as low as 0.095 μg/L (24-hour LC50 value).^6^ Other fish species, such as brook trout (*Salvelinus fontinalis*; BT) and freshwater rainbow trout (*Oncorhynchus mykiss*; RBT), also showed acute mortalities at concentrations of 6PPD-Q of 0.59 μg/L (24-hour LC50) and 1.00 μg/L (96-hour LC50), respectively.^7^ Other acutely sensitive fishes may include white-spotted char (*Salvelinus leucomaenis*), and anadromous steelhead trout (*Oncorhynchus mykiss*), although these fishes were exposed to urban roadway runoff as opposed to isolated 6PPD-Q exposure.^8-9^ Toxicity of 6PPD-Q appears to be highly species-specific, and many of the tested fish species to date do not show any abnormal physiological responses or acute mortality, even after exposure to much greater, non-environmentally relevant concentrations.^7-9,11^ For example, Arctic char (*Salvelinus alpinus*; AC) and white sturgeon (*Acipenser transmontanus*; WS) were tolerant to concentrations greater than 14.2 and 12.7 μg/L, respectively.^7^ Other tolerant species include Atlantic salmon (*Salmo salar*; AS) and brown trout (*Salmo trutta*).^11^ Interestingly, sensitivity cannot be explained by phylogeny alone, as evolutionarily closely related salmonid species, such as AC and BT differ greatly in their responses.^7^

Interspecific differences in response to acute exposure could be attributed to a variety of factors. One explanation could be differences in the molecular targets among different fish species. A recent study using gill cells of RBT suggests that 6PPD-Q results in uncoupling of the electron transport chain, causing increased oxygen consumption, in line with the often-observed symptoms of gasping and spiraling.^7,12^ This study reported effects that were similar to results observed with a known uncoupler, carbonyl cyanide-p-trifluoromethoxyphenylhydrazone.^12^ However, mitochondrial respiration is well-conserved among different species of fishes, and thus, an alternative explanation or other contributing factors may be plausible in explaining the species-specific toxicity of 6PPD-Q.^13^ Another explanation could involve differences in toxicokinetic properties among species. For example, the abundance of a chemical in the body might be influenced by relative capacities to biotransform the parent chemical into a metabolite.^14^ The same might help explain differences in 6PPD-Q sensitivity in fishes.

This work highlights the characterization of the biotransformation of 6PPD-Q as a possible factor contributing to the stark differences in sensitivity among fish species. In accordance with a previous article that hypothesized the plausibility of a toxicokinetic mechanism as a driver of interspecific differences in responses, we hypothesize that lesser metabolic capacity to rapidly detoxify 6PPD-Q may be related to increased sensitivity.^12^ Specifically, this study aimed to identify and characterize potential differences in detoxification of 6PPD-Q across diverse species of sensitive and tolerant fishes through analysis of bile samples from previously published and newly performed experiments using liquid chromatography and high-resolution mass spectrometry (LC-MS).

## MATERIALS AND METHODS

### Chemicals and Reagents

Deuterium-labeled 6PPD-Q-d5 standards were obtained either from HPC Standards GmbH (Borsdorf, Germany) for the Chinook salmon (*Oncorhynchus tshawytscha*; ChS) and CS exposures, from Trent University (Peterborough, ON, Canada; synthesized by Dr. Pirrung) for AS exposures, or from Toronto Research Chemicals (Toronto, ON, Canada) for all other species. Stock solutions for exposure of 6PPD-Q to fish were prepared using ethanol (EtOH) for ChS and CS or dimethyl sulfoxide (DMSO) for all other tested species to achieve a final solvent concentration of 0.0007 % (v/v) EtOH (ChS and CS) and 0.001% - 0.04% (v/v) DMSO during exposures. Analytical standard solutions of native and mass-labeled 6PPD-Q were prepared in absolute EtOH (ChS and CS) or LC-MS-grade methanol (MeOH, all other species). EtOH was obtained from Fisher Scientific (USA). LC-MS grade MeOH and water for bile dilution samples were obtained from Thermo Scientific (Canada) and used as the LC solvents. Extractions were conducted with methanol (LC-MS grade). No standards were available for the identified metabolites of 6PPD-Q due to the novelty of the discovered compounds.

### Fish Source and Culture

RBT and WS were obtained from Lyndon Hatcheries (ON, Canada) and the Nechako White Sturgeon Conservation Centre (Vanderhoof, BC, Canada), respectively, and cultured under standard conditions at the Aquatic Toxicology Research Facility (ATRF) at the University of Saskatchewan (SK, Canada). BT were from Allison Creek Trout Hatchery (Coleman, AB, Canada) and were housed in the Aquatic Research Facility (ARF) at the University of Lethbridge (Lethbridge, AB, Canada). Details of fish source and culture for BT, RBT, and WS can be found in Brinkmann et al. 2022.^7^

Westslope cutthroat trout (*Oncorhynchus clarkii lewisi*; WCT) were obtained from Allison Creek Trout Hatchery (Coleman, AB). WCT were housed for eight months in 600-L insulated fiberglass tanks in the Aquatic Research Facility (ARF) at the University of Lethbridge, AB, Canada, prior to the 6PPD-Q exposure. The photoperiod was 14 h light: 10 h dark. Water flow in the tank was 30 L/min. Water quality was assessed weekly during the holding period. Average values were as follows: DO = 98% (9.8 mg/L), pH = 7.90, ammonia = 0.01 mg/L, nitrite = 0.002 mg/L, nitrate = 3.0 mg/L. Fish were fed EWOS trout feed (Cargill, Canada) at 1% of their body mass per day. Studies were approved by the University of Lethbridge Animal Welfare Committee (Protocol 2111).

CS and ChS were from Voights Creek Hatchery (Orting, WA, USA). Fish were housed in 1,000-L fiberglass circular tanks at the Aquatic Toxicology Lab, Washington State University Puyallup Research and Extension Centre, Puyallup, WA, USA, for 205 days (CS) and 235 days (ChS) days prior to exposure. The photoperiod was 12 h light: 12 h dark. Water flow was recirculated using an aquaculture system with reconstituted water treated by biofiltration and UV. Conductivity and alkalinity are maintained by an auto-dosing system with stocks of Instant Ocean brand sea salt and bicarbonate. The water flow rate was 14.2 L/min in each rearing tank. Water quality was assessed daily (temperature, DO, pH, conductivity, and ammonia) or biweekly (nitrate, nitrite, phosphate, alkalinity) during the holding period. Average values were: DO ≥ 8mg/L, pH = 7.5, conductivity = 800-1,000 μS/cm. Fish were fed 2 % of body mass three times a week with BioOregon BioVita fry feed. Studies were approved by Washington State University’s Institutional Animal Care and Use Committee.

AS were from in-house brood stock at the Huntsman Marine Science Centre and held for ∼1 year (Parr) or ∼2.5 years (Post-Smolt and Post-Smolt Returns) in 7,500-L round tanks prior to exposure. The parr and post-smolt returns were exposed in freshwater, while the post-smolt exposures were conducted in natural seawater. The photoperiod was 10 h light: 14 h dark. Water quality was assessed during the holding period. Average (± SD) values were as follows: temperature: 11.7 ± 0.19 °C (Post-Smolt Returns), 11.4 ± 0.43 °C (Post-Smolt), 13.3 ± 0.27 °C (Parr); DO: 108.9 ± 7.56% (68.5 ± 0.93 mg/L) (Post-Smolt Returns), 122.7 ± 5.42% (11.1 ± 0.61 mg/L) (Post-Smolt), 104.8 ± 2.68% (9.4 ± 0.77 mg/L) (Parr). Fish were fed with a commercial salmonid diet purchased from Skretting. All holding and care for fish were conducted according to the Department of Fisheries and Oceans Animal Care Committee protocol AUP 22-26.

### Exposure Experiments

BT, RBT, and WS were exposed to concentrations of 6PPD-Q ranging from 0.11 to 4.35, 0.09 to 5.33, and 12.7 μg/L, respectively, under static renewal conditions (50% water change every 24 h).

Detailed descriptions of exposure conditions, study design and analytical and quality control procedures for BT, RBT, and WS can be found in Brinkmann et al. 2022.^7^

WCT exposures were performed in 150-L inert glass fiber Krescel tanks under non-renewal conditions with a 2.5 L/min turnover. WCT were exposed to either a solvent control (0.01% v/v DMSO) or a nominal 6PPD-Q concentration of 10 μg/L in 0.01% v/v DMSO. There were four replicate tanks per treatment with four fish in each. Two tanks from control and treatment were sampled at 4 h and 24 h.

Tanks were continuously aerated, and water was recirculated over the course of the exposure. Fish were acclimated to tanks for three days prior to exposure. Fish were fasted for 24 h prior to exposure. A photoperiod of 14 h light: 10 h dark was used. Average (±SD) water quality parameters were as follows: temperature =10.1 ± 0.5 °C; DO = 100.3 ± 8.2%.

ChS and CS exposures were performed in 212-L glass aquaria with silicone seals under static conditions. ChS and CS were exposed to either a solvent control (0.0007% EtOH) or a LC-MS verified 6PPD-Q concentration of 0.04, 0.07, or 0.08 μg/L (CS) and 2.5 μg/L (ChS). There were two replicates per concentration with eight fish each. Tanks were aerated with two glass air diffusers per tank, and no water changes were necessary for the 24-h exposure. Fish were acclimated to tanks for eight hours prior to exposure at 10 °C. Fish were fasted for ∼24 hours prior to exposure. A photoperiod of 12 h light: 12 h dark was used. Average (± SD) water quality parameters were: DO = 9.30 ± 0.24 mg/L; pH = 7.81 ± 0.09; conductivity = 1,124 ± 15.86 μS/cm; temperature = 10.25 ± 0.31 °C. Water samples were collected for analytical confirmation of 6PPD-Q concentrations at time points 0, 3, and 24 hours. Water samples were immediately spiked at 0.1 μg/L with 6PPD-Q-d5 and stored at 4 °C until they were analyzed.

AS exposures were performed in 212-L steel drums with a plastic bag during exposure under static conditions. AS were exposed to either a solvent control (0.04 % v/v DMSO) or a nominal 6PPD-Q concentration of 3, 10, 30, or 100 μg/L. There were three replicates per treatment with five fish each (Post-Smolt and Post-Smolt Returns) or ten fish each (Parr). All tanks from control and treatment were sampled at 24 h of exposure. Tanks were aggressively aerated, and water was not changed over the course of the exposure. Fish were acclimated in the test vessel for 0.5-1 hour at ∼11.5 °C (Post-Smolt and Post-Smolt Returns) or ∼13 °C (Parr) prior to dosing. Fish were fasted for three days (Post-Smolt and Post-Smolt Returns) or two days (Parr) prior to exposure. A photoperiod of 10 h light: 14 h dark was used.

Average (± SD) water quality measurements were as follows: Pre-exposure: Salinity = 33 PSU (Post-Smolt); pH = 7.8 (Post-Smolt Returns), 8.25 (Post-Smolt), 6.88 (Parr); Hardness = 13 mg/L (Post-Smolt Returns), 13 mg/L (Parr); Alkalinity = 20 (Post-Smolt Returns), 24 mg/L (Parr). Post-exposure parameters: Salinity = 32.0 - 33.0 PSU (Post-Smolt); pH = 6.87 - 7.81 (Post-Smolt Returns), 7.65 – 7.96 (Post-Smolt), 7.16 – 7.29 (Parr); Hardness = 12 mg/L (Post-Smolt Returns), 12 mg/L (Parr); Alkalinity = 20 - 28 mg/L (Parr); Ammonia = 0.56 – 1.08 mg/L (Post-Smolt Returns), 0.16 – 0.83 mg/L (Post-Smolt) and 0.08 – 0.33 mg/L (Parr).

### Biological Sampling

WCT were euthanized using 1 g/L Tricaine methanesufonate (TMS) at 4 and 24 h and gallbladders removed and placed in 2-mL cryovials.. CS and ChS were euthanized by percussive stunning followed by pithing. ChS and CS gallbladders were excised and pooled for each replicate into glass vials. Bile was extracted by removing gallbladder tissues with jeweller forceps within collection vials and immediately flash frozen in liquid nitrogen. Bile from ChS was collected at the end of the 24 h exposure period, while for CS, bile was collected from control and 6PPD-Q exposed fish at 8, 9, 11, 19, or 23 h for the few fish that became moribund at those times or 24 h for the remaining fish.

In the AS trials, surviving parr, post-smolt, and post-smolt returns were euthanized with 400 mg/L of TMS at the end of the 24 h exposure. AS bile was extracted from gallbladders using a 1-mL syringe and aliquoted directly into cryovial tubes and subsequently flash-frozen using liquid nitrogen. WS bile was collected from control and 6PPD-Q exposed fish at 96 h of exposure. RBT bile was expressed from gallbladders directly into cryovials that were subsequently flash-frozen in liquid nitrogen. Bile was collected from control and 6PPD-Q exposed fish at ∼12, 16, 60 h for moribund fish, and 96 h for the remaining fish. BT bile was extracted and subsequently frozen at -20°C. Bile was collected from control and 6PPD-Q exposed fish at ∼3, 5, 6, 10, 14 h for moribund fish and 24 h for the remaining fish.

### Sample Processing

Bile samples generated by collaborators were shipped to the University of Saskatchewan on dry ice and stored at -80 °C until further analysis. Samples were thawed at 4 °C or on ice and subsequently aliquoted into MeOH (LC-MS grade) for extraction with a final dilution percentage of 1% for BT, RBT, WS, WCT, 10% for CS, 5% for ChS and 1, 2 or 5% for AS (**Table S2**). After dilution, bile samples were set aside for a minimum of 12 hours at 4 °C for protein. Larger volumes of diluted bile (500 μL) were subsequently filtered through 0.2 μm PTFE filters obtained from Chromatographic Specialties (Canada), and plastic two-piece 3-mL syringes from Thermo Scientific (Canada). Smaller diluted bile samples for CS (30 μL) were centrifuged for 15 minutes at 2 °C and 3,381 ×*g*, and the remaining supernatant was extracted for chemical analyses.

### Instrument Analysis

#### Non-Targeted Analysis Method

Details on the separation of analytes using ultra-high performance liquid chromatography (UPLC) coupled with an Exactive HF Orbitrap high-resolution mass spectrometer (HESI) ion source (Thermo Scientific) are provided in **Table S3**. The following parameters for non-target analysis of bile extracts were used to acquire the data-dependent MS2 (ddMS2) positive mode scans: sheath gas flow = 35, aux gas flow = 10, sweep gas flow = 1, aux gas heater temperature = 300 °C, spray voltage = 4.00 kV, S-lens RF = 60.0, capillary temperature = 350 °C. A Full MS method used scan settings of resolution = 120,000, positive ion, AGC target = 500,000, maximum injection time = 100 ms, Full MS scan range of 70-1,000 m/z. The ddMS2 component of the method used the following scan settings: resolution = 30,000, positive ion, AGC target = 100,000, maximum injection time = 100 ms, loop count = 5, MSX count 2, TopN = 10, with an isolation window of 2.0 m/z and collision energy of 15, 30 and 45 kV.

#### Parallel Reaction Monitoring Method

Details on the separation of analytes using ultra-high performance liquid chromatography (UPLC) coupled with an Exactive HF Orbitrap high-resolution mass spectrometer (HESI) ion source (Thermo Scientific) are provided in **Table S3**. The following parameters for targeted analysis of bile extracts were used to acquire the data-dependent MS2 (ddMS2) positive mode scans: sheath gas flow = 35, aux gas flow = 10, sweep gas flow = 1, aux gas heater temperature = 300 °C, spray voltage = 4.00 kV, S-lens RF = 60.0, capillary temperature = 350 °C. A Full MS/parallel reaction monitoring (PRM) method used the following scan settings: resolution = 30,000, positive ion, AGC target = 2e5, maximum injection time = 100 ms, Full MS scan range of 100.0-1,000 m/z and PRM isolation window of 4.0 m/z. Inclusion list ions, collision energies, and retention times are provided in **Table S4**.

Semi-quantification was achieved using peak areas quantified in Qual Browser Xcalibur (Thermo Scientific) from data generated by the Q-Exactive HF Orbitrap. No standards were available for the discovered compounds, and therefore, the data reported should be considered semi-quantitative. Although full quantitation cannot be achieved without a standard curve, abundance of the metabolites relative to each other are precise and reliable.

### Data Analysis and Statistics

A list of suspected biotransformation products of 6PPD-Q was generated *in silico* using the online tool BioTransformer 3.0 (https://biotransformer.ca/). Data from non-targeted screening were observed in Thermo Scientific FreeStyle software to determine precursor and fragment ions of potential biliary compounds and compared against these *in silico* predictions. To elucidate the most likely molecular position of chemical biotransformation, we used the tool CFM-ID (https://cfmid.wishartlab.com/) to predict MS/MS fragments of 6PPD-Q biotransformation products predicted by BioTransformer 3.0. Once fragments were identified and included within the PRM inclusion list, Thermo Excalibur Qual Browser was used to determine metabolite peak areas in chromatograms in conjunction with fragments within the mass spectra. Peak areas for each metabolite were normalized to 1% bile dilutions if dilutions had more than 1% bile, and then exported into GraphPad Prism 9 software (La Jolla, FL, USA) for subsequent analyses.

Data transformations were necessary to ensure datasets met the requirements of normal distribution (Shapiro-Wilk’s test) and homoscedasticity (Bartlett’s test) prior to statistical testing. A logarithmic function was used to transform phase I data (**Figure 1 A, C**) and phase II data (**Figure 1 B, D**). A one-way ANOVA and subsequent post-hoc Tukey’s test was conducted for these datasets. No transformations could successfully normalize the data in (**Figure 1 E, F**). Therefore, a Kruskal-Wallis with Dunn’s post-hoc test was performed on these respective datasets. A principal component analysis (PCA) was performed using GraphPad Prism 9 software to elucidate the extent to which exposure concentration, species sensitivity (LC50), and the abundance of both metabolites are related to each principal component (**Figure 3**).

**Figure 1:**
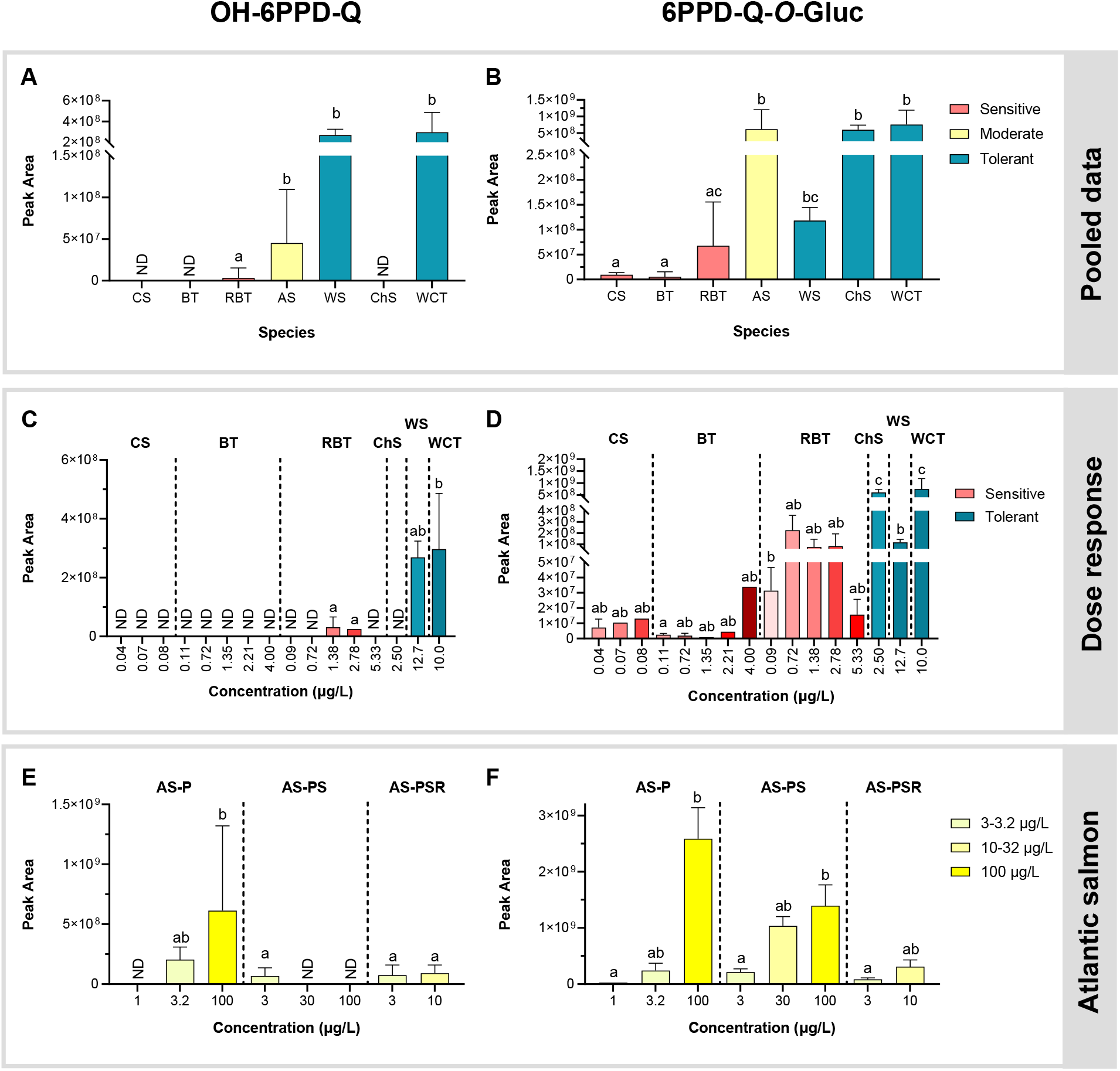
Peak areas indicating the relative abundance of phase I OH-6PPD-Q (**A, C, E**) and phase II 6PPD-Q-*O*-Gluc (**B, D, F**) metabolites in bile samples from various species of fishes. Data in panels **A** and **B** were compiled according to the closest possible match in measured 6PPD-Q exposure concentration, with the intention of highlighting relationships with species sensitivity. Data in panels **C** and **D** depict the available data across species and measured exposure concentrations of 6PPD-Q, except for Atlantic salmon (AS), while panels **E** and **F** show comparisons of the abundance of phase I and II metabolites across different AS life stages and exposure concentrations of 6PPD-Q. Data from coho salmon (CS) and chinook salmon (ChS) were generated from pooled bile samples, while data from AS, brook trout (BT), rainbow trout (RBT), white sturgeon (WS), and westslope cutthroat trout (WCT) were obtained from individual fish. Suffixes in AS samples are defined as follows: P – Parr, PS – Post-Smolt, PSR – Post-Smolt Returns. Groups marked with different letters were statistically different from one another. *ND – not detected*.

## RESULTS AND DISCUSSION

### Toxicity of 6PPD-Q to Test Species

The responses of BT, RBT, and WS to exposure with 6PPD-Q were described previously (**Table S1**).^7^ AS Parr and later life (Post-Smolt and Post-Smolt Returns) stages displayed tolerance to very high nominal concentrations of 6PPD-Q (100 μg/L), while fry displayed sensitivity to exposure (**Table S1**). CS displayed few mortalities (8/53) at the exposure concentrations (≤ 8 μg/L), while ChS displayed tolerance to 6PPD-Q exposure to verified concentrations of 2.5 μg/L (**Table S1**). WCT were insensitive to a nominal 6PPD-Q concentration of 10 μg/L (**Table S1**). No mortalities were seen in WCT, WS, RBT, BT AS, CS, and ChS exposures in control treatments.

### Analytical Identification of 6PPD-Q Metabolites

Using a full-MS ddMS2 non-targeted analytical workflow, we discovered two suspected metabolites of 6PPD-Q: (1) mono-hydroxy-6PPD-Q (OH-6PPD-Q) and (2) 6PPD-*O*-glucuronide (6PPD-Q-*O*-Gluc) (**Figures S1-S4**). OH-6PPD-Q (m/z 315.17) showed a mass-to-charge ratio of parent 6PPD-Q (m/z 299.17) + 16 (**Figure S4**). The 6PPD-Q-*O*-Gluc metabolite showed an m/z of OH-6PPD-Q + 176, which is characteristic of glucuronide conjugation (**Figure S3**). Fragments for both metabolites appeared to be present in their respective mass spectra. No other biotransformation products predicted by BioTransformer 3.0 beyond the two described here could be confirmed in the tested species. The CFM-ID MS/MS fragment predictor was used to output potential fragments for different OH-6PPD-Q and 6PPD-Q-*O*-Gluc isomers, whereby only those modified at the aryl moiety contained fragments which matched the observed spectra (**Figure S4**). Based on these comparisons, it appeared most likely that the initial hydroxylation occurs at the phenyl ring adjacent to the quinone rather than the alkyl moiety (**Figure 2**).

**Figure 2:**
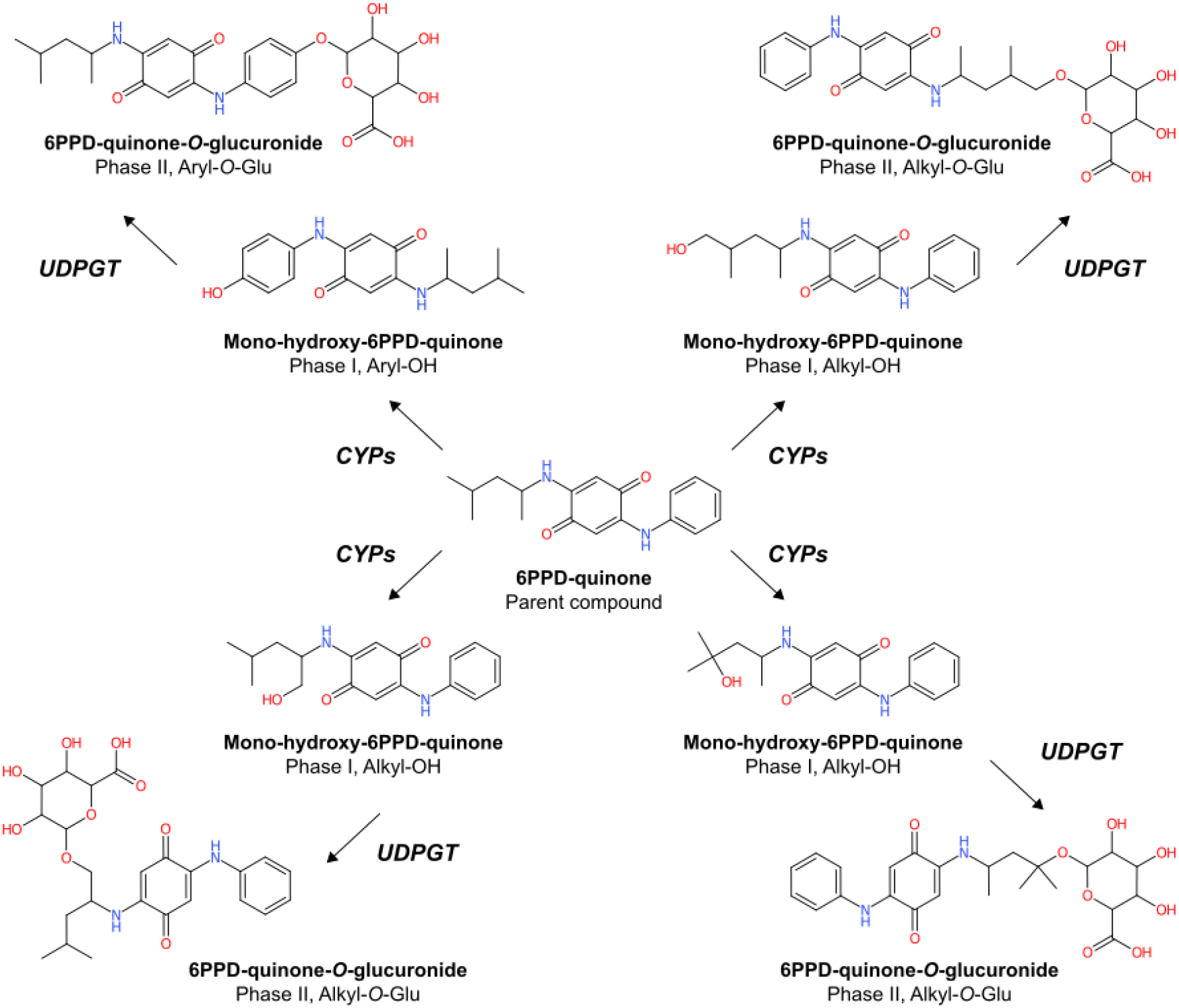
*In silico* predictions for the biotransformation of 6PPD-quinone into a phase I mono-hydroxy-metabolite and a subsequent phase II *O*-glucuronide metabolite. Comparison of MS2 spectra with those predicted by CFM-ID suggested the presence of aryl-substituted (top left) instead of alkyl-substituted metabolites.

We recognize that this information is only of preliminary nature, and follow-up investigations using NMR and other methods are required to fully characterize these tentatively identified metabolites.

### Analytical Verification of 6PPD-Q Metabolites

Based on the findings of the non-target ddMS2 method, we developed a parallel reaction monitoring (PRM) method using the characteristic transitions summarized in **Table S4**. The PRM method successfully eliminated irrelevant noise and resulted in increased sensitivity and resolution for the targeted biotransformation products. Instrumental chemical analyses of bile samples using the newly developed PRM methods revealed significant differences in OH-6PPD-Q (phase I) and 6PPD-Q-*O*-Gluc (phase II) metabolite abundance, which varied between species (**Figure 1 A, B**), as well as within species at varying exposure concentrations of 6PPD-Q exposure (**Figure 1 C-F**). Specific comparisons are discussed and further examined in the following sections.

### Interspecific Differences Based on Pooled Data

Both OH-6PPD-Q and 6PPD-Q-*O*-Gluc were not detectable in bile liquid of control treatments of any of the investigated species (data not shown). This is an important observation that confirms that there is no endogenous formation of these two metabolites when not exposed to 6PPD-Q. Similarly, OH-6PPD-Q was not detected in the bile of CS, BT, and ChS. The average abundance of OH-6PPD-Q in the bile liquid of RBT (3.37 × 10^7^ arbitrary units, au) was significantly lower than that in the bile liquid of AS (4.52 × 10^7^ au), WS (2.69 × 10^8^ au), and WCT (2.97 × 10^8^ au). With the exception of ChS, the abundance of OH-6PPD-Q was smallest in the most sensitive species and increased with decreasing sensitivity (**Figure 1A**). Similarly, the abundance of 6PPD-Q-*O*-Gluc was lowest in the three sensitive species (CS, BT, and RBT) and increased with decreasing sensitivity (**Figure 1B**). However, it should be acknowledged that the only sensitive life stage of AS (fry) was not represented in the bile dataset due to their small size (see below), and hence, further studies are required to characterize metabolite abundance in the more sensitive life stage of AS. Overall, these findings suggest that metabolite profiles might be useful to distinguish species that are tolerant or sensitive to 6PPD-Q exposure.

Factors that could contribute to the observed differences in abundance of the two metabolites are manifold, including not only differences in metabolic capacity but also potential impacts of temperature on passive absorption, distribution, and elimination.^15,16^ However, temperatures of exposure were similar for all tested species: most were tested at 10 °C, with some tested at 12 ± 1 °C. Therefore, it is improbable that differential rates of passive transport of 6PPD-Q into the gills contributed to metabolite abundance study. Although temperature is an unlikely contributing factor, absorption rates of 6PPD-Q may still have played a major role in metabolite abundance. Metabolite levels in WCT 24 h samples were either 4.1

(OH-6PPD-Q) or 5.8 (6PPD-Q-*O*-Gluc) times larger than in 4 h samples, indicating the acute duration of 6PPD-Q exposure influenced initial metabolite abundance. Although, the total duration of exposure typically followed a 24-hour time course for non-moribund fishes, moribund fishes were sampled earlier, thereby yielding decreased metabolite abundance. Little is known about the absorption, distribution, metabolism and excretion (ADME) properties of 6PPD-Q. Preferential distribution of 6PPD-Q throughout the body is not yet discerned and the rate of 6PPD-Q uptake and biotransformation into or *via* the liver has not yet been published. Other ADME factors, such as active transport into the liver or rapid elimination rates, might have contributed toward the measured abundance of metabolites. However, further studies are needed to fully elucidate the contribution of these potential factors.

The most likely contributing factor could be differences in metabolic capacity to biotransform 6PPD-Q between sensitive and insensitive species. In this context, lower 6PPD-Q metabolite abundance in sensitive versus insensitive species might be attributed to two potential factors. First, 6PPD-Q was shown to uncouple mitochondrial respiration in RBT gill cells,^12^ which is known to lead to a depletion of ATP pools and in turn, a reduction in overall metabolic activity. This could have directly impacted metabolite abundance by limiting the clearance of 6PPD-Q. However, mitochondrial respiration is well conserved among all fishes and unlikely to differ sufficiently between species to explain differences in metabolite abundances. The second and more plausible explanation is that basal expression of biotransformation enzymes responsible for catalyzing phase I and/or phase II reactions might be substantially lower in sensitive species compared to insensitive species. This hypothesis requires further investigation, as the specific enzymes and/or isoforms catalyzing the first hydroxylation reaction of 6PPD-Q, e.g., cytochrome P450-dependent monooxygenases (CYPs), and the UDP glucuronosyltransferase (UDPGT) isoforms responsible for glucuronidation, have not been identified yet. Based on our findings, however, we hypothesize tolerant species and life stages of fish can efficiently remove 6PPD-Q from their circulation prior to the onset of acute lethality.

Generally, the levels of OH-6PPD-Q were about one order of magnitude lower compared to those of 6PPD-Q-*O*-Gluc across all samples (**Figure 1**). This observation is not uncommon for biotransformation processes since expression levels and activities of phase II enzymes in vertebrates are normally several orders of magnitude greater than those of phase I enzymes, thereby efficiently facilitating the removal of a variety of phase I biotransformation products. In a previous study, this can be seen when comparing the activities of ethoxyresorufin *O*-deethylase (EROD), which is catalyzed by CYP1A, versus those of UDPGT and glutathione *S*-transferase (GST), in RBT hepatocytes.^17^ In this study, UDPGT activity was, on average, 50-fold greater than that of EROD, and GST activity was 40,000-fold greater.^17^ As a result, levels of phase II biotransformation products are often considerably greater than those of phase I products. For example, the levels of *O*-glucuronides of the polycyclic aromatic hydrocarbon benzo[*a*]pyrene (BaP) were a factor of roughly 100 greater than those of OH-BaP in early-life stages of fathead minnows (*Pimephales promelas*) following exposure to BaP.^17,18^

### Intraspecific Differences Based on Exposure Concentration and Duration

When comparing OH-6PPD-Q abundance across species (except for AS) and exposure concentrations (**Figure 1C**), most samples showed levels below the sensitivity threshold of the analytical instrument. In addition, for the few samples that showed measurable levels OH-6PPD-Q, there were no statistically significant differences across concentrations within each species.

6PPD-Q-*O*-Gluc levels showed concentration-dependent trends among some of the investigated species (**Figure 1D**). For example, the abundance of 6PPD-*O*-Gluc in CS exposed to 0.07 and 0.08 μg/L 6PPD-Q were a factor of 1.4 and 1.8, respectively, greater than in those exposed at 0.04 μg/L. However, these differences were not significant due to the necessity to pool CS samples due to their low volume. Similarly, in BT exposed to 0.11 μg/L and 4.0 μg/L, the abundance of 6PPD-*O*-Gluc was a factor of 13.6 greater in the 4.0 μg/L treatment. However, this difference was not significant due to the low number of surviving fish at the highest exposure concentration.

Interestingly, we observed an inverse U-shaped concentration response in RBT. Here, 6PPD-Q-*O*-Gluc abundance appeared to increase between 0.09 to 0.72 μg/L, followed by a decrease between 0.72 to 5.33 μg/L (**Figure 1D**). This pattern of lower phase II metabolite abundance is likely a result of the earlier onset of mortalities with increasing concentrations and a consequent decrease in exposure duration of analyzed fish. The 96-hour LC50 value for RBT was determined to be 1.00 μg/L, which corresponds to the cut-off where the decrease in phase II metabolite abundance began (**Figure 1D**).^7^ To study the impact of exposure time further, both time and concentration-dependence of metabolite abundance should be the subject of future research.

### Differences in Atlantic Salmon Across Life Stages and Concentrations

Significant differences were detected in OH-6PPD-Q abundance in AS bile across the various life stages tested (**Figure 1E**). Specifically, there was a concentration-dependent increase in OH-6PPD-Q abundance at the Parr stage, which was significantly greater at the 100 μg/L exposure concentration (6.13 × 10^8^ au) compared to the other life stages and concentrations, ranging from non-detected to 2.05 × 10^8^ au (**Figure 1E**). Similarly, Parr showed the greatest abundance of 6PPD-Q-*O*-Gluc in bile liquid, followed by Post-smolt and Post-Smolt Returns (**Figure 1F**). However, while there were concentration-dependent increasing trends in metabolite abundance at the same life stage, these differences were only statistically significant in 6PPD-Q-*O*-Gluc data. The observed concentration responses are likely more conclusive compared to BT and RBT since these life stages were not sensitive, and all fish were exposed for the same duration. Sampling of the only sensitive life stage of AS, fry, was not possible due to their small size. Therefore, we could not determine conclusively whether there are significant differences in metabolite abundance across life stages that could help explain the observed life stage differences in sensitivity. However, the three life stages of Parr, Post-Smolt Returns, and Post-Smolt showed similarly low sensitivity and similar levels of 6PPD-Q-*O*-Gluc. It is expected that the sensitive fry life stage of AS would have a lower presence of metabolites due to the trends described above. In addition to this, earlier life stages of fish tend to have lower CYP expression than their older counterparts.^19^ It is possible that CYPs are responsible for the initial detoxification step of 6PPD-Q; however, this needs further elucidation, as no published studies to date have examined biotransformation pathways of 6PPD-Q in fish.

### Multi-Variate Analysis for LC50, Exposure Concentration and Metabolite Abundance

The principal component analysis (PCA) revealed that principal component 1 (PC1) explained 45.75% of the variance in the data, while PC1 and PC2 together explained 73.56% of the variance within the data (**Figure 3**). Scores for insensitive species clustered in the same quadrant, typically as an individual group or otherwise clustered as one insensitive group (**Figure 3**), while sensitive species scores closely clustered as one collective group (**Figure 3**). The scores for AS appeared to cluster in three separate groups, with no obvious relation to the other sensitive or insensitive clusters (**Figure 3**). In PC1, the small negative exposure duration loading pointed in the opposite direction of all other large positive loadings (**Figure 3**). In PC2, the small positive duration, large positive OH-6PPD-Q and large positive LC50 loadings all point in the opposite direction of the two other loadings, that is the small negative 6PPD-Q-*O*-Gluc and large negative exposure concentration loadings (**Figure 3**). Based on these results, it appears that samples from sensitive and insensitive species, as well as AS were distinct from another.

**Figure 3:**
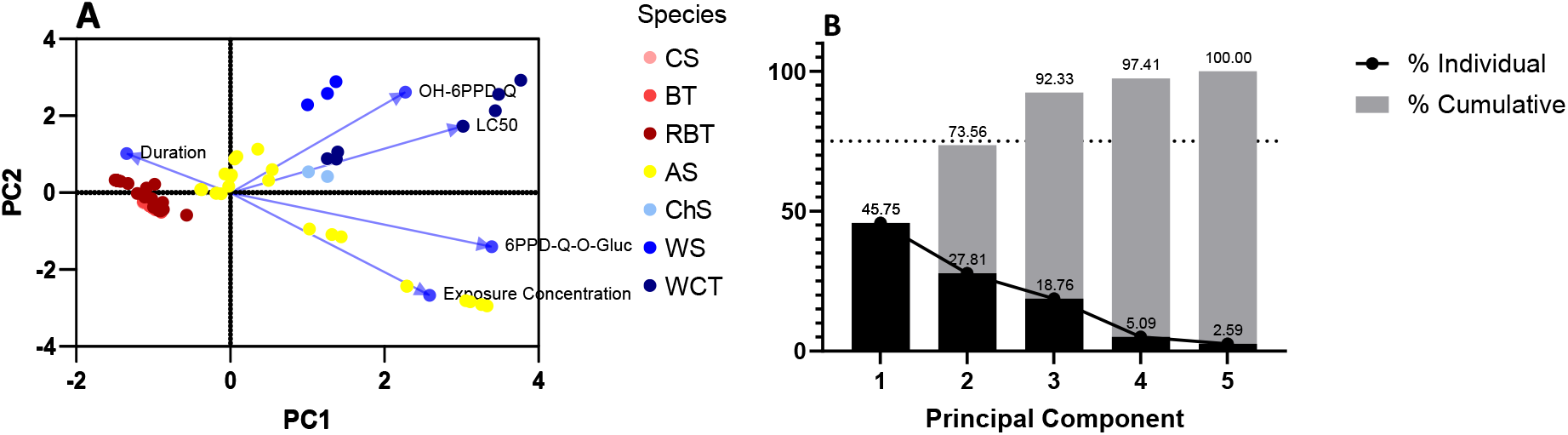
Principal component analysis (PCA) biplot (**A**) and proportions of variance (**B**) for exposure concentration, duration of exposure, LC50, OH-6PPD-Q, and 6PPD-Q-*O*-Gluc. The dotted line in B indicates the typical cut-off of 75% for total variance used in PCA.

More specifically, OH-6PPD-Q abundance and LC50 were highly correlated (R = +0.706) throughout the dataset. Therefore, it appears phase I detoxification of 6PPD-Q is the major driver for distinguishing sensitive from insensitive species. Although the correlation matrix depicts a correlation between 6PPD-Q-*O*-Gluc and LC50 (R = +0.487), the strength of correlation is weaker than its phase I counterpart, OH-6PPD-Q, which further supports the notion that the phase I reaction is primarily responsible for detoxification of 6PPD-Q. While 6PPD-Q-*O*-Gluc and exposure concentration also appear to be strongly correlated (R = +0.800), the correlation between OH-6PPD-Q and exposure concentration did not appear to be strong (R = -0.058). Therefore, exposure concentration may have played a role in 6PPD-Q-*O*-Gluc abundance but is unlikely for OH-6PPD-Q abundance. All correlations with exposure duration showed very low values (ranging from R = -0.132 to +0.009), therefore it is unlikely exposure duration played a major role in metabolite abundance in the present dataset. Based on these results, it appears that 6PPD-Q-*O*-Gluc abundance was driven mostly by exposure concentration, and could therefore be considered a biomarker of exposure, while the abundance of OH-6PPD-Q was driven mostly by the sensitivity of the species (LC50) and could therefore be an important indicator of species sensitivity.

### Outlook and Future Research Needs

This study is the first to test the hypothesis that differential detoxification rates of 6PPD-Q could contribute toward the highly species-specific acute lethality of this compound. However, several studies are required to evaluate this potential mechanism further. The variability of duration and concentrations of exposure makes comparisons difficult within this study. Further testing of bile samples from the same sublethal exposure concentrations for the same study duration and life stage of fish might help elucidate this further. To confirm the major route of detoxification is in the liver, *in vitro* substrate depletion assays exposing hepatocytes of different species to 6PPD-Q may provide a more comprehensive comparison for the relative rates of hepatic 6PPD-Q biotransformation. Additionally, co-exposure with different enzyme-specific inhibitors might help pinpoint the identity of responsible biotransformation enzyme(s) and/or isoform(s). Lastly, the chemical synthesis and further characterization of neat chemical standards for each of the tentatively identified metabolites should be attempted, thereby increasing confidence and permitting quantitative determination of these metabolites in fish samples.

Based on the findings described above, detection of 6PPD-Q-*O*-Gluc in the bile of field-exposed fish might be a useful biomarker of exposure to 6PPD-Q. A sublethal exposure of 6PPD-Q to fish *in vivo*, should contribute significantly to determining the residence time of each metabolite and further elucidate the feasibility of 6PPD-Q-*O*-Gluc as a biomarker of exposure. Similar to the determination of biliary metabolites of polyaromatic hydrocarbons in fish,^20^ this method might enable improved environmental monitoring of exposure of wild-caught or caged fish and has the potential to tremendously benefit the environmental risk assessment of 6PPD-Q.

## Acknowledgments

This project was supported partially by a financial contribution from Fisheries and Oceans Canada. Additional funding was provided to M.B., M.H., and S.W. through the Discovery Grants program of the Natural Sciences and Engineering Research Council of Canada (NSERC). Instrumentation used for chemical analyses within this research was obtained with funding from Western Economic Diversification Canada (WED) and the Canadian Foundation for Innovation (CFI). M.B. is currently a faculty member of the Global Water Futures (GWF) program, which received funds from the Canada First Research Excellence Funds (CFREF). S.W. and M.H. were supported by the Canada Research Chairs Program. The authors acknowledge the animal care support of Zoë Henrikson and Dale Jefferson (ATRF) and Holly Shepherd and Mamun Shamsuddin (ARF). Analytical support was provided by Xiaowen Ji and Jenna Cantin. The authors also acknowledge the general assistance provided by Katherine Raes, Lucy Kapronczai, Evan Kohlman, and Phillip Ankley.

## Notes

### Competing Interest Statement

The authors have declared no competing interest.

## References

(1) Product-Chemical Profile for Motor Vehicle Tires Containing 6PPD - Final Version. (n.d.). from https://dtsc.ca.gov/wp-content/uploads/sites/31/2022/05/6PPD-in-Tires-Priority-Product-Profile_FINAL-VERSION_accessible.pdf

(2) Peter (2022). Where Does Rubber For Tires Come From? (n.d.). Retrieved February 5, 2023 Product-Chemical Profile for Motor Vehicle Tires Containing 6PPD - Final Version. (n.d.). from https://dtsc.ca.gov/wp-content/uploads/sites/31/2022/05/6PPD-in-Tires-Priority-Product-Profile_FINAL-VERSION_accessible.pdf

(3) Classification of antioxidants used in rubber tires. (n.d.). Henan Xuannuo Chemicals Co. Ltd. Retrieved February 5, 2023, from https://www.xnadditive.com/info/classiff

(4) Challis, J. K., Popick, H., Prajapati, S., Harder, P., Giesy, J. P., McPhedran, K., & Brinkmann, M. (2021). Occurrences of Tire Rubber-Derived Contaminants in Cold-Climate Urban Runoff. Environmental Science & Technology Letters, 8(11), 961–967. https://doi.org/10.1021/acs.estlett.1c00682

(5) Johannessen, C., Helm, P., Lashuk, B., Yargeau, V., & Metcalfe, C. D. (2022). The Tire Wear Compounds 6PPD-Quinone and 1,3-Diphenylguanidine in an Urban Watershed. Archives of Environmental Contamination and Toxicology, 82(2), 171–179. https://doi.org/10.1007/s00244-021-00878-4

(5) Tian, Z., Zhao, H., Peter, K. T., Gonzalez, M., Wetzel, J., Wu, C., Hu, X., Prat, J., Mudrock, E., Hettinger, R., Cortina, A. E., Biswas, R. G., Kock, F. V. C., Soong, R., Jenne, A., Du, B., Hou, F., He, H., Lundeen, R., … Kolodziej, E. P. (2021). A ubiquitous tire rubber–derived chemical induces acute mortality in coho salmon. Science & Technology Letters, 9(9), 733–738. https://doi.org/10.1021/acs.estlett.2c00467

(6) Tian, Z., Gonzalez, M., Rideout, C. A., Zhao, H. N., Hu, X., Wetzel, J., Mudrock, E., James, C. A., McIntyre, J. K., & Kolodziej, E. P. (2022). 6PPD-Quinone: Revised Toxicity Assessment and Quantification with a Commercial Standard. Environmental Science & Technology Letters, 9(2), 140–146. https://doi.org/10.1021/acs.estlett.1c00910

(7) Brinkmann, M., Montgomery, D., Selinger, S., Miller, J. G. P., Stock, E., Alcaraz, A. J., Challis, J. K., Weber, L., Janz, D., Hecker, M., & Wiseman, S. (2022). Acute toxicity of the Tire Rubber-Derived Chemical 6PPD-quinone to Four Fishes of Commercial, Cultural, and Ecological Importance. Environmental Science & Technology Letters, acs.estlett.2c00050. https://doi.org/10.1021/acs.estlett.2c00050

(8) Hiki, K., & Yamamoto, H. (2022). The Tire-Derived Chemical 6PPD-quinone Is Lethally Toxic to the White-Spotted Char Salvelinus leucomaenis pluvius but Not to Two Other Salmonid pecies. Environmental Science & Technology Letters, 9(12), 1050–1055. https://doi.org/10.1021/acs.estlett.2c00683

(9) French, B. F., Baldwin, D. H., Cameron, J., Prat, J., King, K., Davis, J. W., McIntyre, J. K., & Scholz, N. L. (2022). Urban Roadway Runoff Is Lethal to Juvenile Coho, Steelhead, and Chinook Salmonids, But Not Congeneric Sockeye. Environmental Science, 371(6525), 185–189. https://doi.org/10.1126/science.abd6951

(10) Lo, B. P., Marlatt, V. L., Liao, X., Reger, S., Gallilee, C., & Brown, T. M. (n.d.). Acute toxicity of 6PPD-quinone to early life stage juvenile Chinook (Oncorhynchus tshawytscha) and coho (Oncorhynchus kisutch) salmon. Environmental Toxicology and Chemistry, n/a(n/a). https://doi.org/10.1002/etc.5568

(11) Foldvik, A., Kryuchkov, F., Sandodden, R., & Uhlig, S. (2022). Acute Toxicity Testing of the Tire Rubber–Derived Chemical 6PPD-quinone on Atlantic Salmon (Salmo salar) and Brown Trout (Salmo trutta). Environmental Toxicology and Chemistry, 41(12), 3041–3045. https://doi.org/10.1002/etc.5487

(12) Mahoney, H., da Silva Junior, F. C., Roberts, C., Schultz, M., Ji, X., Alcaraz, A. J., Montgomery, D., Selinger, S., Challis, J. K., Giesy, J. P., Weber, L., Janz, D., Wiseman, S., Hecker, M., & Brinkmann, M (2022). Exposure to the Tire Rubber-Derived Contaminant 6PPD-Quinone Causes Mitochondrial Dysfunction In Vitro. Environmental Science & Technology Letters, 9(9), 765–771. https://doi.org/10.1021/acs.estlett.2c00431

(13) Irene, V., & Sazanov, L. A. (2022). The assembly, regulation and function of the mitochondrial respiratory chain. Nature Reviews.Molecular Cell Biology, 23(2), 141–161. doi:https://doi.org/10.1038/s41580-021-00415-0

(14) Ribalta, C., Sanchez-Hernandez, J. C., & Sole, M. (2015). Hepatic biotransformation and antioxidant enzyme activities in Mediterranean fish from different habitat depths. Science of The Total Environment, 532, 176–183. https://doi.org/10.1016/j.scitotenv.2015.06.001

(15) Zhernenkov, M., Bolmatov, D., Soloviov, D., Zhernenkov, K., Toperverg, B. P., Cunsolo, A., Bosak, A., & Cai, Y. Q. (2016). Revealing the mechanism of passive transport in lipid bilayers via phonon-mediated nanometre-scale density fluctuations. Nature Communications, 7(1), Article 1. https://doi.org/10.1038/ncomms11575

(16) Xu, H., Feng, C., Cao, Y., Lu, Y., Xi, J., Ji, J., Lu, D., Zhang, X.-Y., & Luan, Y. (2019). Distribution of the parent compound and its metabolites in serum, urine, and feces of mice administered 2,2′,4,4′-tetrabromodiphenyl ether. Chemosphere, 225, 217–225. https://doi.org/10.1016/j.chemosphere.2019.03.030

(17) Fay, K. A., Fitzsimmons, P. N., Hoffman, A. D., & Nichols, J. W. (2017). Comparison of Trout Hepatocytes and Liver S9 Fractions as In Vitro Models for Predicting Hepatic Clearance in Fish. Environmental Toxicology and Chemistry, 36(2), 463–471. https://doi.org/10.1002/etc.3572

(18) Grimard, C., Mangold-Döring, A., Schmitz, M., Alharbi, H., Jones, P. D., Giesy, J. P., Hecker, M., & Brinkmann, M. (2020). In vitro-in vivo and cross-life stage extrapolation of uptake and biotransformation of benzo[a]pyrene in the fathead minnow (Pimephales promelas). Aquatic Toxicology, 228, 105616. https://doi.org/10.1016/j.aquatox.2020.105616

(19) Saad, M., Cavanaugh, K., Verbueken, E., Pype, C., Casteleyn, C., Van Ginneken, C., & Van Cruchten, S. (2016). Xenobiotic metabolism in the zebrafish: A review of the spatiotemporal distribution, modulation and activity of Cytochrome P450 families 1 to 3. Journal of Toxicological Sciences, 41(1), 1–11. https://doi.org/10.2131/jts.41.1

(20) Kammann, U., Akcha, F., Budzinski, H., Burgeot, T., Gubbins, M. J., Lang, T., Le Menach, K., Vethaak, A. D., & Hylland, K. (2017). PAH metabolites in fish bile: From the Seine estuary to Iceland. Marine Environmental Research, 124, 41–45. https://doi.org/10.1016/j.marenvres.2016.02.014

