## Supporting Information for "Toxicokinetic Characterization of the Inter-Species Differences in 6PPD-Quinone Toxicity Across Seven Fish Species: Metabolite Identification and Semi-Quantification"

Total pages: 8

Tables: 4

Figures 4

*Corresponding author:

**Markus Brinkmann, PhD**

Toxicology Centre, University of Saskatchewan

44 Campus Drive

Saskatoon, SK, S7N 5B3 Canada

**Table S1.** Proportion of samples where fish displayed classic symptoms of exposure to 6PPD-Q, such as gasping, etc. prior to death and their respective LC50 (where applicable). Sensitivity data were collected from Tian et al. 2022, Brinkmann et al. 2022, and unpublished data from Huntsman Marine Science Centre and the Wiseman lab.

| **Species** | **LC50 (24 hour)** | **Sample Comments** |
|---|---|---|
| Coho salmon (*Oncorhynchus kisutch*) | 0.096 µg/L | Bile was extracted from fish exposed to 0.045, 0.066, and 0.084 µg/L for 24 h with few (8/53) mortalities |
| Brook trout (*Salvelinus fontinalis*) | 0.59 µg/L | Bile was extracted from fish exposed to 0.11, 1.35, and 2.21, 4.00 µg/L for 24 h with majority of mortalities occurring in the first 12 h |
| Rainbow trout (*Oncorhynchus mykiss*) | 1.96 µg/L | Bile was extracted from fish exposed to 0.09, 0.72, 1.38, 2.78 and 5.33 µg/L for 96 h with majority of mortalities occurring within the first 24 h |
| Atlantic salmon (*Salmo salar*) | Early life stages are sensitive. However, all bile tested was extracted from later life stages tolerant to 6PPD-Q | Bile was extracted from fishes exposed to nominal concentrations of 3, 3.2, 10, 30, 32 and 100 µg/L for 24 h |

| | | |
|---|---|---|
| White sturgeon (*Acipenser transmontanus*) | Tolerant to ~12.7 µg/L | Bile was extracted from fishes exposed to 12.7 µg/L for 96 h |
| Chinook salmon (*Oncorhynchus tshawytscha*) | Tolerant to 2.5 µg/L | Bile was extracted from fishes exposed to 2.5 µg/L for 24 h |
| Westslope cutthroat trout (*Oncorhynchus clarkii lewisi*) | Tolerant to nominal ~10 µg/L | Bile was extracted from fishes exposed to nominal 10 µg/L for 4 to 24 h |

**Table S2.** Extracted percentages of bile diluted in LC-MS grade methanol for each tested species. Bile samples were frozen during isolation, thawed prior to extraction, and analyzed using Thermo Q-Exactive HRMS analysis.

| **Species** | **Percent Bile** | **Percent Methanol** |
|---|---|---|
| Coho salmon (*Oncorhynchus kisutch*) | 10% | 90% |
| Brook trout (*Salvelinus fontinalis*) | 1% | 99% |
| Rainbow trout (*Oncorhynchus mykiss*) | 1% | 99% |
| Atlantic salmon (*Salmo salar*) | 1, 2, 5% | 99, 98, 95% |
| White sturgeon (*Acipenser transmontanus*) | 1% | 99% |
| Chinook salmon (*Oncorhynchus tshawytscha*) | 5% | 95% |
| Westslope cutthroat trout (*Oncorhynchus clarkii lewisi*) | 1% | 99% |

**Table S3. Chromatographic conditions for positive mode NTA and PRM methods.** Flow rate = 0.200 mL/min, column temperature 40 °C, solvent A = 95% $H_2O$; 5% MeOH; 0.1% formic acid and B = 100% MeOH; 0.1% formic acid.

| **Time (min)** | **%B** |
|---|---|
| 0.00 | 5.0 |
| 7.50 | 40.0 |
| 15.00 | 100.0 |
| 20.00 | 100.0 |
| 20.01 | 5.0 |
| 25.00 | 5.0 |

**Table S4.** Parent metabolite and fragment ions ($[M+H]^+$), collision energy (CE), and retention time (RT) details for the full-scan parallel reaction monitoring Orbitrap mass spectrometer method.

| **Metabolite** | **Precursor Ion (*m/z*)** | **Fragment Ions (*m/z*)** | **CE** | **RT (min)** |
|---|---|---|---|---|
| Mono-Hydroxy-6PPD-Q | 315.17 | 231.08 | 35 | ~17.3 |
| 6PPD-Q-*O*-Glucuronide | 491.20 | 407.10, 315.17, 231.08 | 35 | ~14.3 |

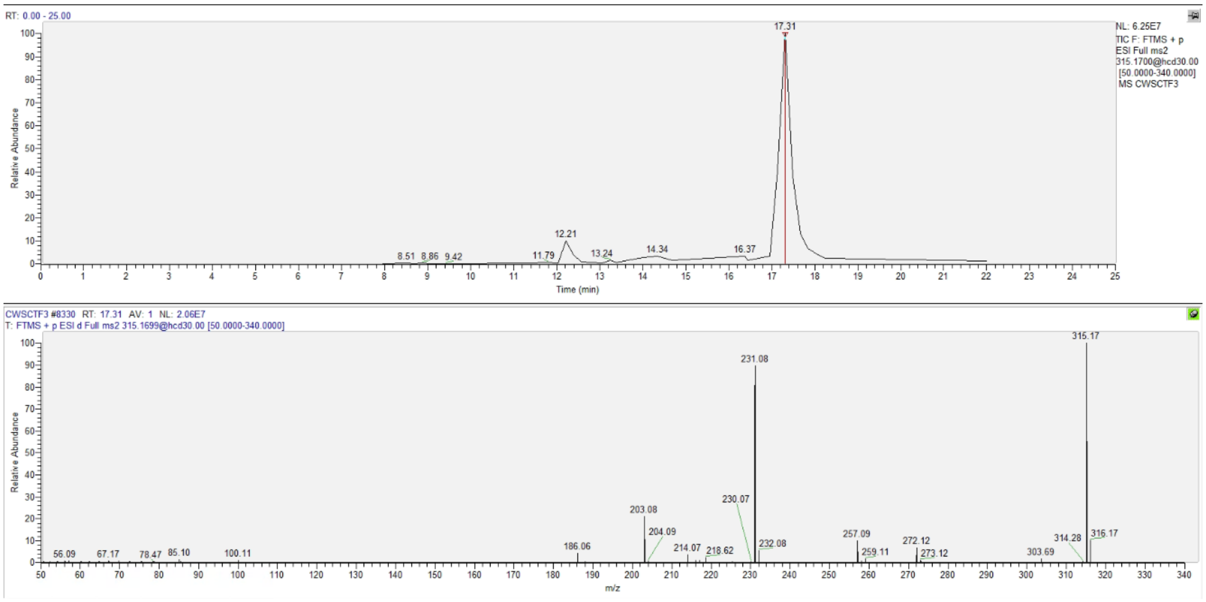


**Figure S1.** Chromatogram and mass spectra for the phase I mono-hydroxy-6PPD-Q metabolite using the non-targeted analyses method (RT = 17.31 min).

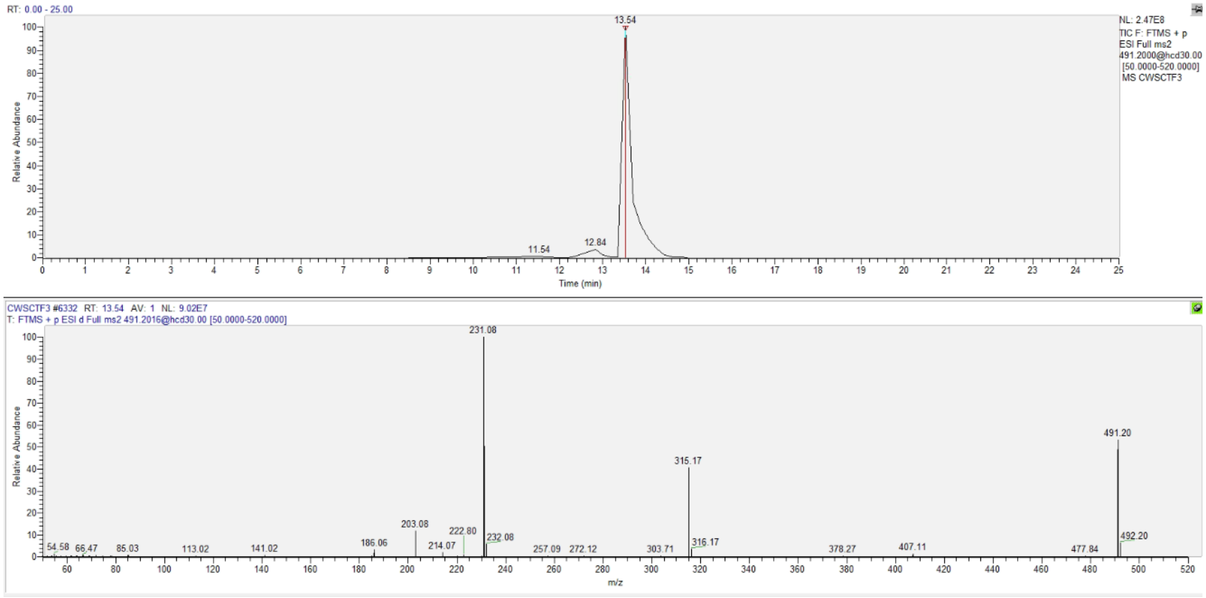


**Figure S2.** Chromatogram and mass spectra for the phase II 6PPD-Q-*O*-glucuronide metabolite using the non-targeted analyses method (RT = 13.54 min).

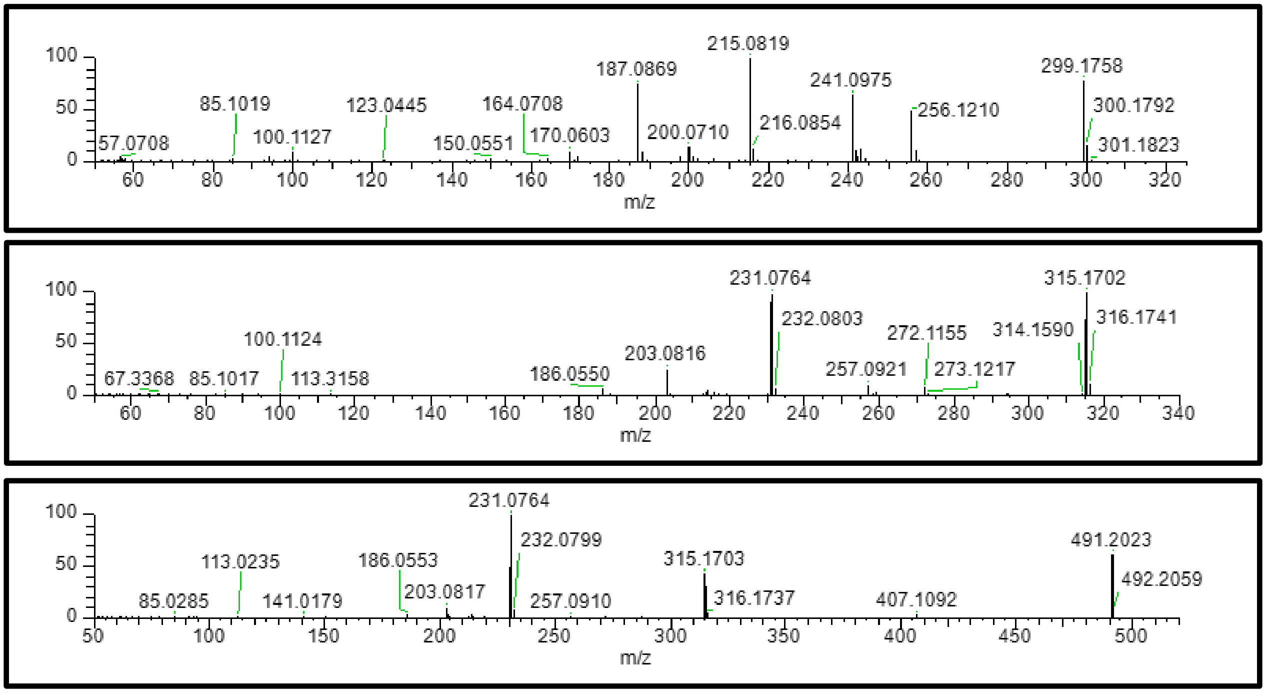


**Figure S3.** Mass spectra for 6PPD-Q (top), the phase I mono-hydroxy-6PPD-Q metabolite (centre) and the phase II 6PPD-Q-*O*-glucuronide metabolite (bottom).

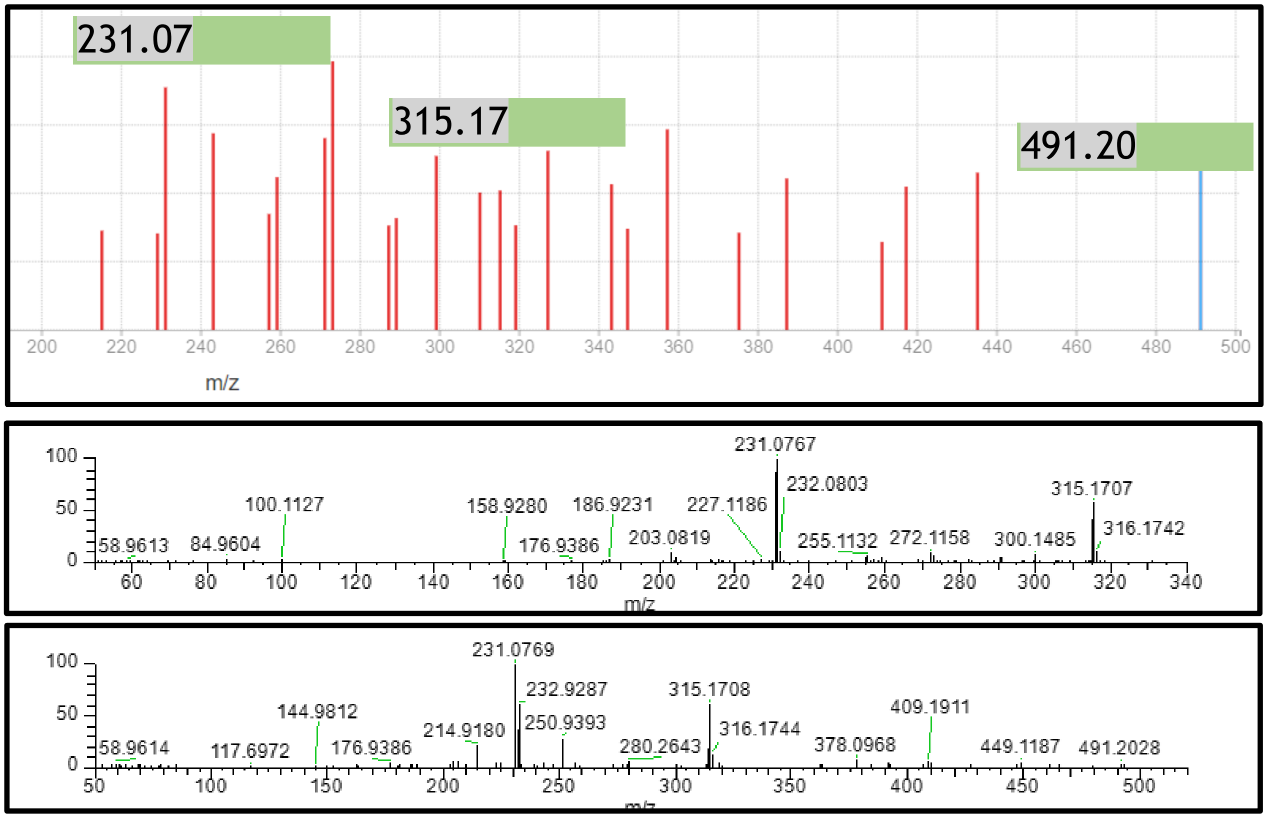


**Figure S4.** CFM-ID output for predicted mass spectra for 6PPD-Q aryl *O*-glucuronide (top) and observed mass spectra for the mono-hydroxy-6PPD-Q metabolite (centre) and 6PPD-Q-*O*-glucuronide metabolite (bottom). The predicted fragment (231.07 *m/z*) is present in both observed mass spectra.
